# Salicylic Acid restricts cell elongation and induces changes of vacuolar morphology and pH

**DOI:** 10.1101/2024.09.06.611645

**Authors:** Jonas Müller, Yvonne König, Sabrina Kaiser, Christian Löfke, Melanie Krebs, David Scheuring

## Abstract

The phytohormone salicylic acid (SA) is a key factor to balance plant defence as well as growth and development. While its role in plant defence has been investigated for decades, regulation of plant growth and development has only come into focus recently. It has been demonstrated that SA application inhibits growth independently of the established Non-expressor of Pathogenesis Related (NPR) receptors. However, the underlying mechanism of this growth inhibition on the cellular level remains largely elusive. Here we show that SA restricts cell elongation and induces changes of vacuolar morphology and pH. Rapidly upon SA application we observe homotypic vacuole fusion and a significant increase in vacuolar pH. These changes seem to be independent of the phytohormone auxin which has been reported to crosstalk with SA. By increasing vacuolar pH, SA directly impacts basic cellular functions such as vesicle trafficking or nutrient storage, leading eventually to cell size restriction and limited growth. Our results demonstrate an NPR-independent mechanism to attenuate growth, potentially allowing free resources to be relocated to withstand environmental stresses.

**Graphical abstract:** 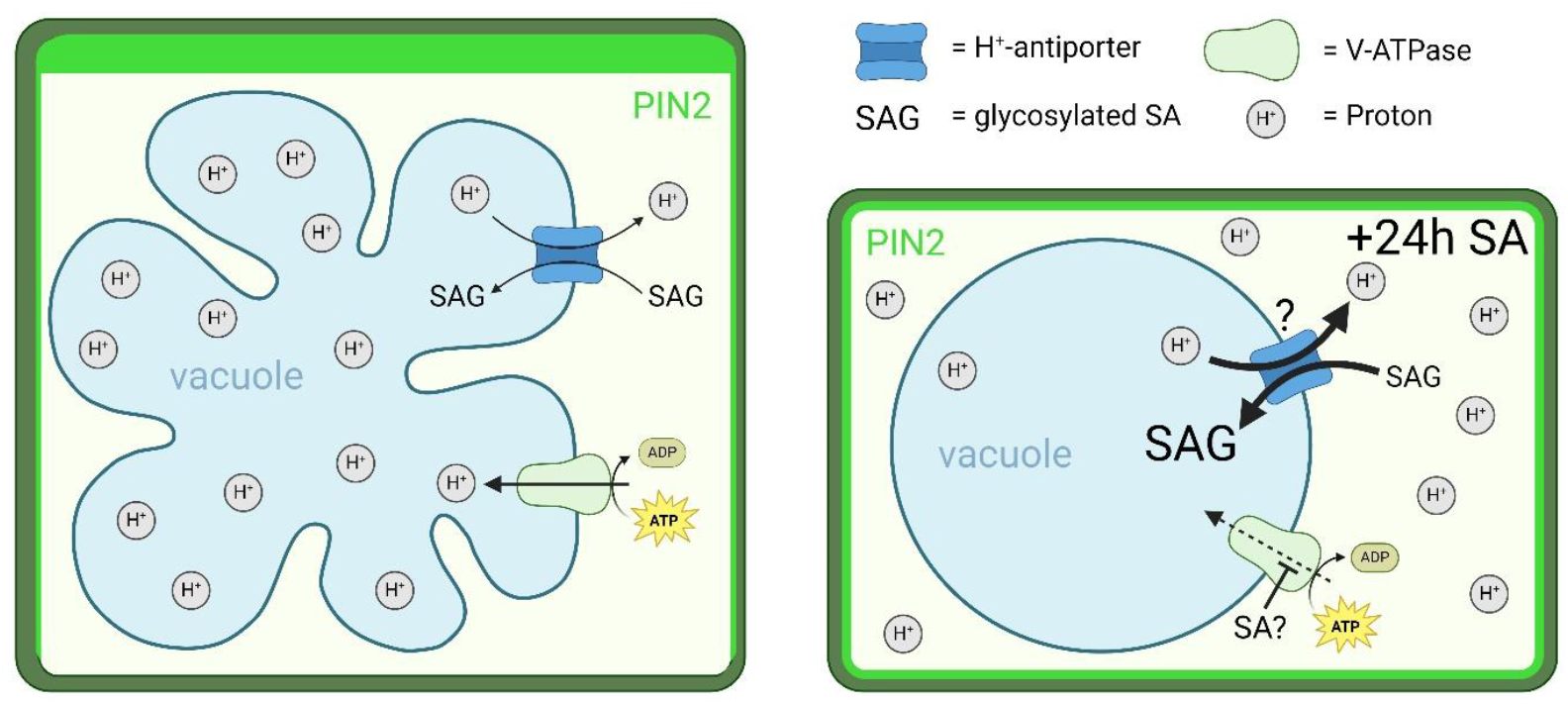

Salicylic Acid (SA) triggers a reduction in cell size and leads to a spherical vacuolar phenotype. This morphological change is accompanied by an increase in vacuolar pH, potentially due to the enhanced activity of a glycosylated SA (SAG) H+-antiporter and/or the inhibition of V-ATPase activity. In addition, SA disrupts the polarity of PIN2 auxin transporters, resulting in their uniform distribution across the cell membrane.

## Introduction

Plants have evolved efficient measures to oppose environmental stresses. Salicylic acid (SA) is one of the key molecules for stress signalling, e.g. in response to pathogen attacks. A rise of endogenous SA levels induces efficient local and systemic defence responses as countermeasure (Loake and Grant, 2007). This comes, however, to the expense of reduced or even inhibited growth. In the last decade, the role of SA in plant defence has been well investigated but the inhibitory effect on plant growth is considerably less understood. Recently, SA function during plant growth and development came into focus (Rivas-San Vicente and Plasencia, 2011). Notably, the signalling cascade mediating SA-induced growth attenuation seems to be independent of its defence-inducing role. It was demonstrated that the *bona fide* SA receptor Nonexpressor of Pathogenesis Related genes 1 (NPR1), the key regulator of plant immune responses, is not responsible for the transmission of growth-restriction when exogenously applied (Tan et al., 2020).

While long distance transport of SA via the phloem to distal tissues to induce systemic acquired resistance (SAR) has been well established (Park et al., 2007), only one intracellular SA transporter has been identified to date (Anfang and Shani, 2021). The MATE family transporter EDS5 was demonstrated to be crucial for SA export from the chloroplast, where it is synthesized, to the cytosol (Serrano et al., 2013). There is, however, biochemical evidence that glycosylated SA is transported into the vacuole for inactivation (Bonnemain et al., 2013). Vacuolar transport was shown to require either an ATP-binding cassette transporter mechanism or an H^+^-antiport mechanism, dependent on the investigated plant species (Dean and Mills, 2004; Dean et al., 2005). The glycosylation step prior vacuole uptake in Arabidopsis is dependent on the small-molecule glucosyltransferases UGT74F1, UGT74F2, and UGT76B1. The latter seems to be central for controlling levels of free and active SA, thereby contributing to balance the plant immune status (Bauer et al., 2021; Saint Paul et al., 2011).

Recently, crosstalk of SA with the phytohormone auxin was demonstrated (Pasternak et al., 2019). SA impacted on auxin synthesis and transport which eventually led to changes in root architecture. Pasternak et al. (2019) showed that low SA concentrations promoted adventitious root emergence and changed the organization of the root meristem, while high SA concentrations (above 250 µM) led to general inhibition of root growth. More recently, also lower SA concentrations were found to attenuate root growth and the reported SA-auxin crosstalk was confirmed: SA binding to A subunits of protein phosphatase 2A (PP2A) led to inhibition of its function and ultimately to changes of the PIN2 auxin transporter (Tan et al., 2020). SA-induced PP2A inhibition causes hyperphosphorylation of PIN2 and loss of its cellular polarity, severely affecting root growth and development. In line with this, endogenous and exogenously applied SA inhibits clathrin-mediated endocytosis and thus affects internalization of plasma membrane proteins such as PINs (Du et al., 2013). However, since PP2A acts on several substrates and *pp2aa1* mutation led to increased SA sensitivity (Tan et al., 2020), it seems likely that other molecular players are involved.

In the past, it was shown that auxin has an inhibiting effect on underground tissues (Evans et al., 1994). Interestingly, auxin has been shown to impact the cells largest organelle – the vacuole directly, leading to smaller vacuoles and restricted cell elongation (Löfke et al., 2015a; Scheuring et al., 2016). In this context, a *space-filling* function was defined for the vacuole: simple inflation and occupation of emerging cellular space allows for rapid cell elongation. Thus, energy consumption for the generation of cytosolic content can be reduced and efficient growth maintained (Scheuring et al., 2016; Krüger and Schumacher, 2018; Dünser et al., 2019; Kaiser and Scheuring, 2020). The auxin-induced reduction in vacuole size is dependent on the actin cytoskeleton (Scheuring et al., 2016; Kaiser et al., 2019) and endocytic trafficking has been demonstrated to promote vacuole enlargement (Dünser et al., 2022). Cell elongation and eventually root growth in turn seems to depend on unconditioned vacuole inflation, leading to the hypothesis that reduction of vacuole size might be a general mechanism to reduce and control growth (Kaiser et al., 2021).

Here, we show that cell size is restricted even by application of moderate SA concentrations. This is accompanied by changes of vacuolar morphology independently of auxin. Furthermore, we demonstrate a rapid increase of vacuole pH, potentially impairing basic cellular functions.

## Results

### SA Specifically Inhibits Root Growth

Recently, it has been demonstrated that SA impacts plant growth and development independently of its *bona fide* receptor NPR1 (Pasternak et al., 2019; Tan et al., 2020). By exogenously applying different SA concentrations on 7-day-old Arabidopsis seedlings, we observed a dosage-dependent root growth reduction Fig. S1A-B). Treatment with 50 µM SA inhibited root growth of the WT and the *npr134* triple mutant to the same extent (Zhang et al., 2006) by 50% (Fig. S1C). Degradation of endogenous SA in the transgenic *nahG* line, expressing a bacterial salicylate hydroxylase gene (Friedrich et al., 1995), led to significantly less root growth inhibition (Fig. S1D). Notably, this growth-inhibiting effect was only found in underground but not aerial tissue: Etiolated seedlings treated with 50 µM SA displayed a 50% reduction in root length but no reduction of hypocotyl size (Fig. 1A and 1B). To confirm hypocotyl responsiveness, we used Concanamycin A (Con. A), an inhibitor of vacuolar H^+^-ATPase, preventing hypocotyl elongation. Application of Con. A showed a significant decrease of hypocotyl length already at 0.2 µM (Fig S1E). Root growth reduction correlated well with SA-induced restriction of maximal size of root epidermal cells in the elongation zone (Fig. 1C and 1D).

**Figure 1:**
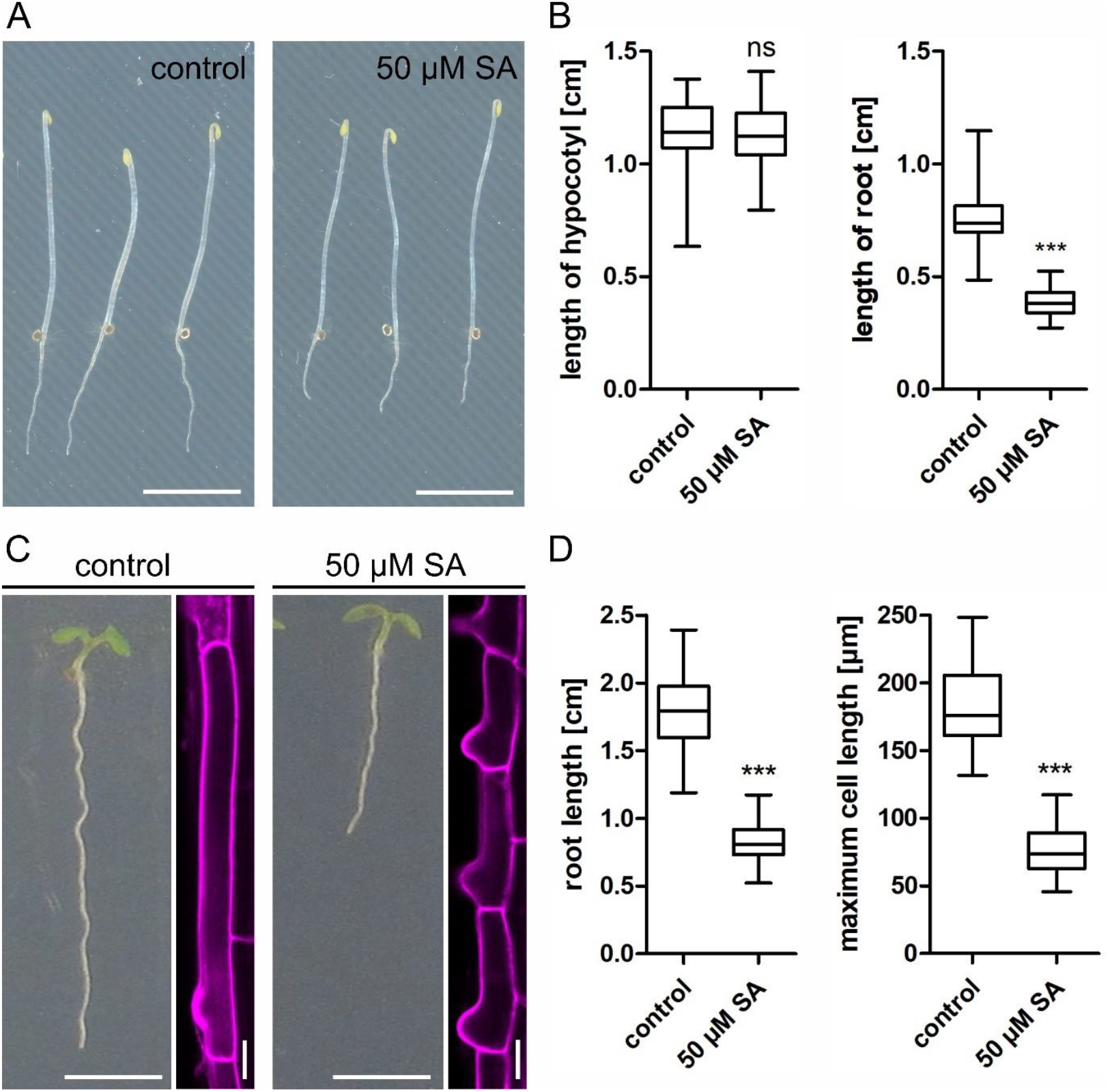
SA reduces root growth and cell size. A, WT Arabidopsis seeds were placed on agar plates and kept in light for 8 hours, then transferred to the dark for an additional 5 days. B, Length of root and hypocotyl after 5 days in the dark (n hypocotyl control = 42, hypocotyl 50 µM SA = 44, root control = 42, root 50 µM SA = 44). Whisker plot with Student’s t-test. *** p≤0.001. Scale bars: 0.5 cm. C, Representative images of Arabidopsis seedlings comparing root length and cell size upon SA treatment. Horizontal scalebars: 0.5 cm; vertical scale bars: 20 µm. D, Quantification of root length and maximum cell elongation in 7-day-old Arabidopsis seedlings (n control = 142, 50 µM SA = 138). Roots were stained with propidium iodide (PI) to highlight cell walls, and fully elongated cells were measured (n control = 28, 50 µM SA = 44). Whisker plot with Student’s t-test. *** p≤0.001.

### SA Reduces Cell Size and Impacts on Auxin by Manipulating PIN2

Reduced root cell size upon auxin treatment has been previously described for (Löfke et al., 2015a). Auxin also affects root and shoot growth differentially, in a concentration dependent manner (Sauer et al., 2013). Consequently, we investigated a potential crosstalk between SA and auxin in epidermal cells of the root meristem. Here, the epidermis is regularly spaced into shorter tricho- and longer atrichoblast cell files and represents a suitable model to study cell size differences (Berger et al., 1998; Löfke et al., 2013). To analyze auxin accumulation upon SA application, we used the semi-quantitative auxin-input reporter R2D2 (Liao et al., 2015). The reporter showed a significant signal down-regulation after 24h SA treatment (Fig. 2A and B), indicating elevated auxin levels. At the same time, we noticed that, in the time frame of 24h, cell size differences of tricho -and atrichoblasts were disappearing (Fig. 2C). In parallel, we observed an SA-induced increase of cell number which was accompanied by an increase in meristem size (Supplemental Fig. S2A, B and C). To determine if this was the result of higher mitotic activity, the B1-type cyclin cell cycle marker *CYCB1;1::GUS* was used (Colón-Carmona et al., 1999). Indeed, upon 24h SA treatment the reporter showed increased activity (Fig. S2D), implying that a higher cell division rate might be responsible for the loss of cell size differences between atrichoblasts and trichoblasts. It has been shown that both cell types exhibit different accumulation of the auxin efflux carrier PIN2 (Löfke et al., 2015b). It was suggested that higher PIN2 concentration observed in atrichoblasts could result in increased auxin export rates and, hence, less cell size restriction. Likewise, lower PIN2 abundance in trichoblasts could slow down auxin export, increasing its inhibitory effect. To test for differences in the distribution of PIN2 in tricho-and atrichoblast cells upon SA treatment we used a functional PIN2-GFP fusion under the endogenous promoter (Abas et al., 2006). Under control conditions, the fluorescence intensity of PIN2-GFP at the apical plasma membrane (PM) in atrichoblasts was higher than that in trichoblast cells as shown before (Löfke et al., 2015b). SA application, however, abolished this difference, resulting in a more uniform PIN2 distribution at the PM of atrichoblasts (Fig. 2E and F). This led to the assumption that auxin is responsible for the reduction in cell size following SA treatment. Since auxin-induced cell size restrictions might be regulated by the size of the plant vacuole (Scheuring et al., 2016; Löfke et al., 2015a), we speculated that SA-induced inhibition of cell size functions similarly. To follow this, we used a PIN2 mutant, *eir1-4*, and the auxin receptor triple mutant *tir1afb2afb3* and tested SA-induced root growth inhibition. For both mutants, no increased resistance against SA could be detected (Fig. 3A and B).

**Figure 2:**
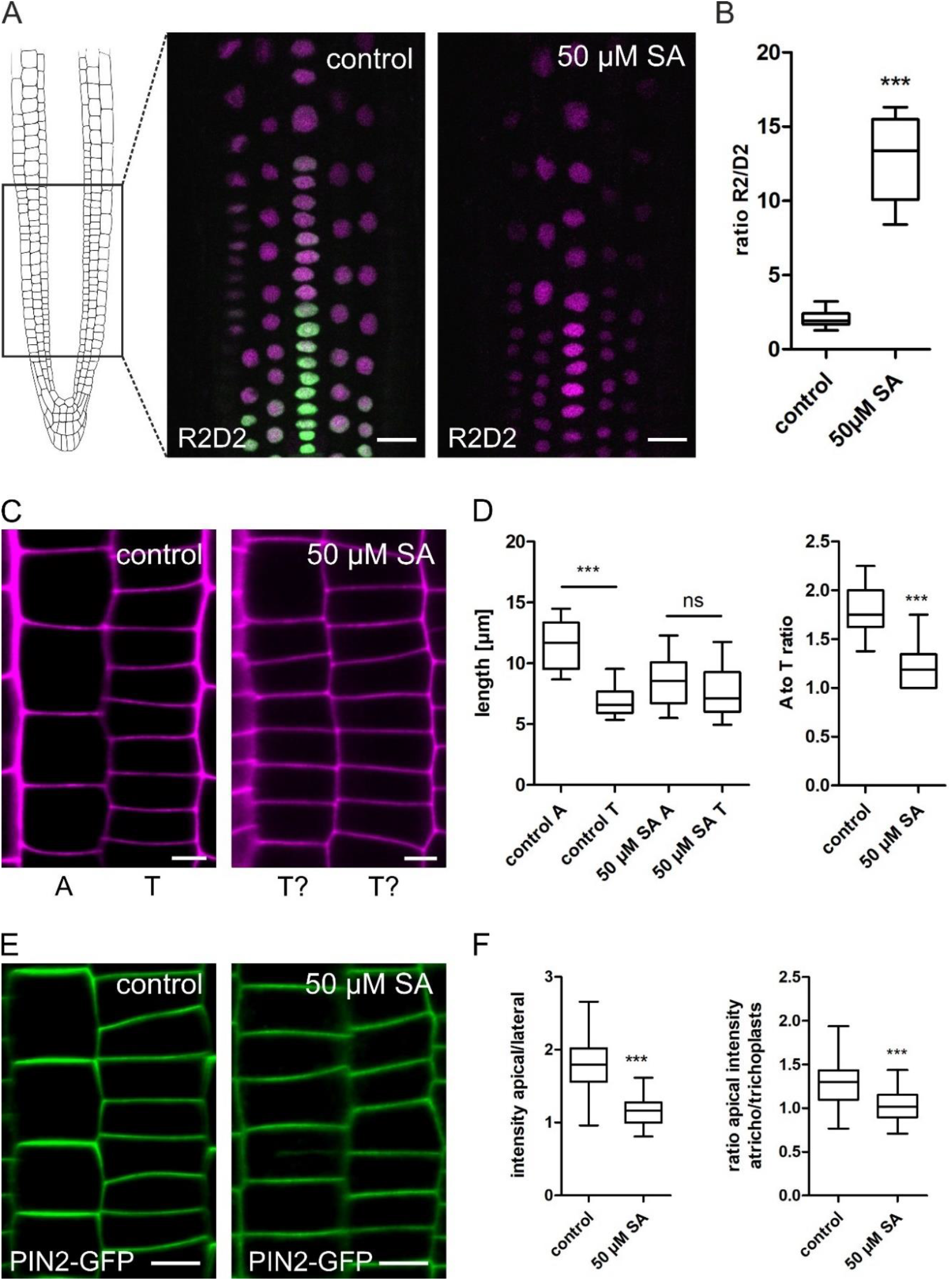
SA reduces cell size and alters PIN2 distribution. A; Area of root for microscopic experiments. Representative images of the rapid auxin-input reporter R2D2 before and after SA treatment. Scale bar: 30 µm. B, Ratio calculation by dividing R2 by D2. Six-day-old R2D2 seedlings were transferred to ½MS+ medium supplemented with 50 µM SA and grown for an additional 24h (n = 19). Whisker plot with Student’s t-test. *** p≤0.001. C, PI-stained epidermal cell files in the root meristem. A = Atrichoblast cell; T = Trichoblast cell. Scale bar: 5 µm. D, Quantification of atrichoblast and trichoblast cell length changes after SA treatment. Six-day-old seedlings were transferred to ½MS+ medium supplemented with 50 µM SA and grown for an additional 24h. The lengths of atrichoblast cells near the transition zone were measured and compared to neighboring trichoblast cells (n = 27). Whisker plot with Student’s t-test. *** p≤0.001. E, Trichoblast and atrichoblast cell files of the PIN2-GFP marker line before and after SA treatment. F, Quantification of the PIN2-GFP intensity ratio of the apical and lateral side of atrichoblast cells. In the further quantification this ratio was divided by the ratio of the apical and lateral side of trichoblast cells. PIN2-GFP seedlings were grown on control plates for 4 days and transferred for 24h on medium supplemented with 50 µM SA (n = 70, n = 10). Whisker plot with Student’s t-test. *** p≤0.001.

**Figure 3:**
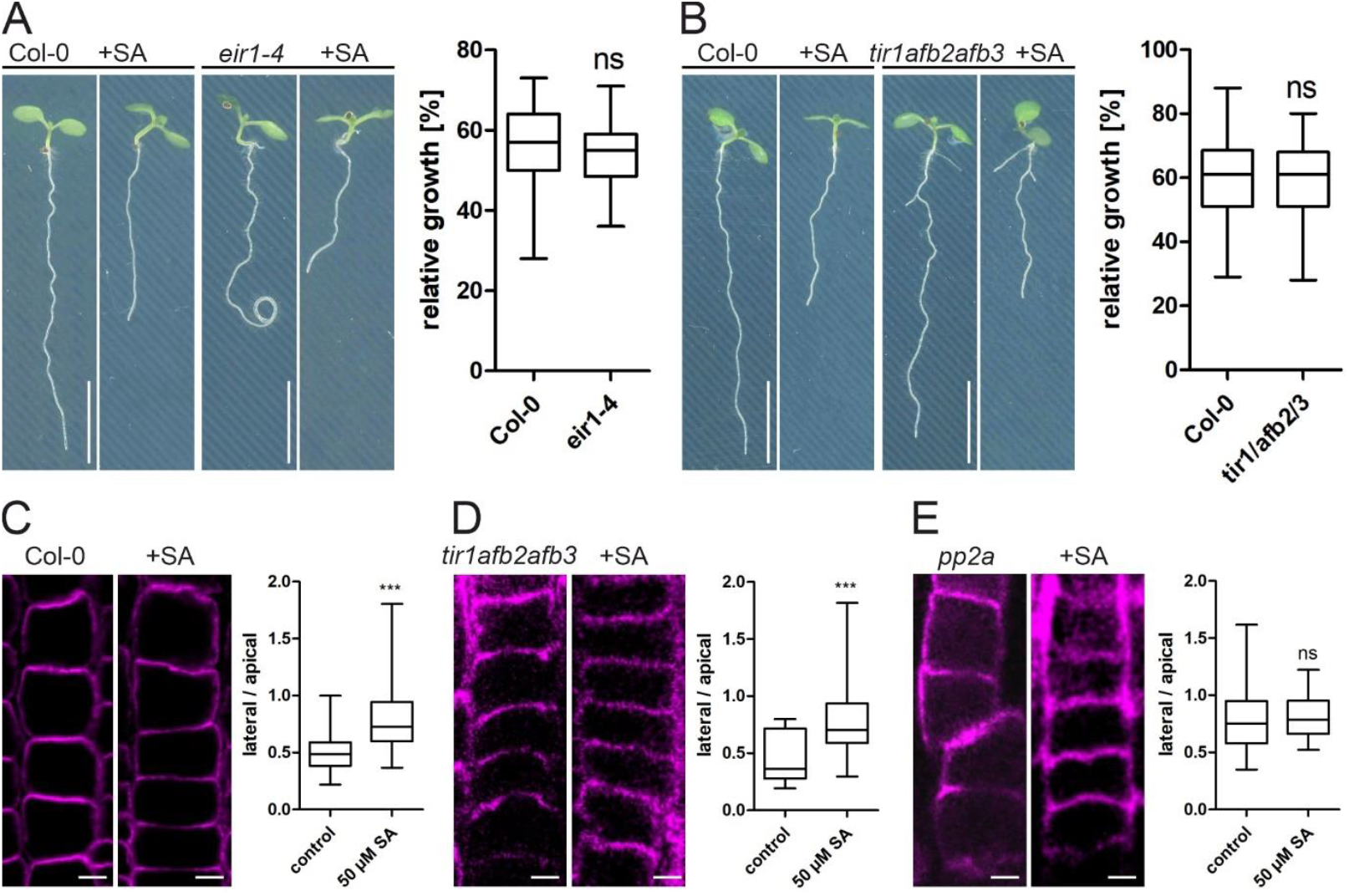
Neither the PIN2 mutant *eir1-4* nor the auxin receptor mutant *tir1afb2afb3* show increased resistance to SA. A-B; Col-0, *eir1-4* and *tir1afb2afb3* seedlings were grown for 7 days on ½MS+ plates and supplemented with 50 µM SA. The relative growth was calculated by dividing each root length of seedlings grown on SA supplemented media with the average root length of its respective control (n *eir1-4* = 29, n *tir1afb2afb3* = 59). Whisker plot with Student’s t-test. ** p≤0.01. Scale bar: 40 mm. C-E, Immunolocalization of PIN2 in Col-0, *tir1afb2afb3* and *pp2a*. Quantification of the intensity ratio of the apical and lateral side of atrichoblast cells. (n Col-0 = 53, *tir1afb2afb3* = 15, *pp2a* = 20). Whisker plot with Student’s t-test. *** p≤0.001.

Using a PIN2 antibody, we also determined PIN2 polarity upon SA application in wt and the *tir1afb2afb3* mutant. Immunolocalization displayed a more uniform PIN2 distribution in SA treated roots similar to the findings with the PIN2-GFP line (Fig. 3C-D). Since SA has been shown to directly bind to PP2A (Tan et al., 2020), we also tested root length and PIN2 polarity in a *pp2a* knockout line: Compared to the wildtype Root growth was more severely affected after SA application and PIN2 polarity was diminished in untreated conditions (Fig. 3E), confirming the reported hypersensitivity of *pp2a* (Tan et al., 2020).

### SA Changes Vacuolar Morphology and Reduces SNARE Abundance

To further investigate the reason for SA-induced root cell growth inhibition, we investigated the effect of SA treatment on vacuolar morphology. Using the *pUBQ10::YFP-VAMP711* as marker to visualize the tonoplast, we observed a change of vacuolar morphology within minutes after SA treatment: vacuoles in epidermal cells of the root meristem displayed homotypic fusion events and became almost entirely spherical within 4h (Movie S1 and S2). To quantifiy these changes, we used the vacuolar morphology index (VMI) to estimate vacuole size (Löfke et al., 2015a). SA application increased the VMI significantly (Fig. 4A and B), while treatments with the SA analogs Acibenzolar-S-methyl (BTH) and 2,6-Dichloroisonicotinic acid (INA) had no effect (Fig. S3A and B). Additionally, treatment of seedlings with 50 µM INA, having a similar pKa value, displayed no root growth inhibition (Fig. S3C). Notably, SA treatment resulted in a completely different vacuolar phenotype compared to auxin-induced changes. In contrast to the SA-induced changes of vacuolar morphology, auxin leads to constricted tubular vacuoles with a decreased VMI (Löfke et al., 2015a; Scheuring et al., 2016). Here, Soluble NSF Attachment Protein Receptor (SNARE) complexes at the vacuole have been shown to be required to conduct auxin-induced constrictions. To test whether disappearance of vacuolar constrictions upon SA treatment involves tonoplast SNAREs, we examined the localization of SYP21-YFP. Interestingly, while exogenous application of auxin increased the fluorescence intensity of tonoplast localised SNAREs including SYP21-YFP (Löfke et al., 2015a), SA treatment decreased SYP21-YFP fluoresence intensity and protein levels in roots (Fig. 4C-E). Induction of homotypic vacuole fusions was further confirmed by using a *net3cnet4anet4b* triple mutant. Together these networked proteins have been shown to redundantly regulate vacuole-vacuole fusions (Kaiser et al., 2023). When treated with SA, root length of *net3cnet4anet4b* was less affected than the corresponding Col-0 wt control (Fig. 4F). Since the mutant roots were already shorter than that of Col-0 in untreated conditions, this resulted in a better relative (Fig. 4G), indicating a partial resistance against SA. Hence, homotypic vacuole fusion seems to be a prerequisite for SA to mediate its inhibitory function.

**Figure 4:**
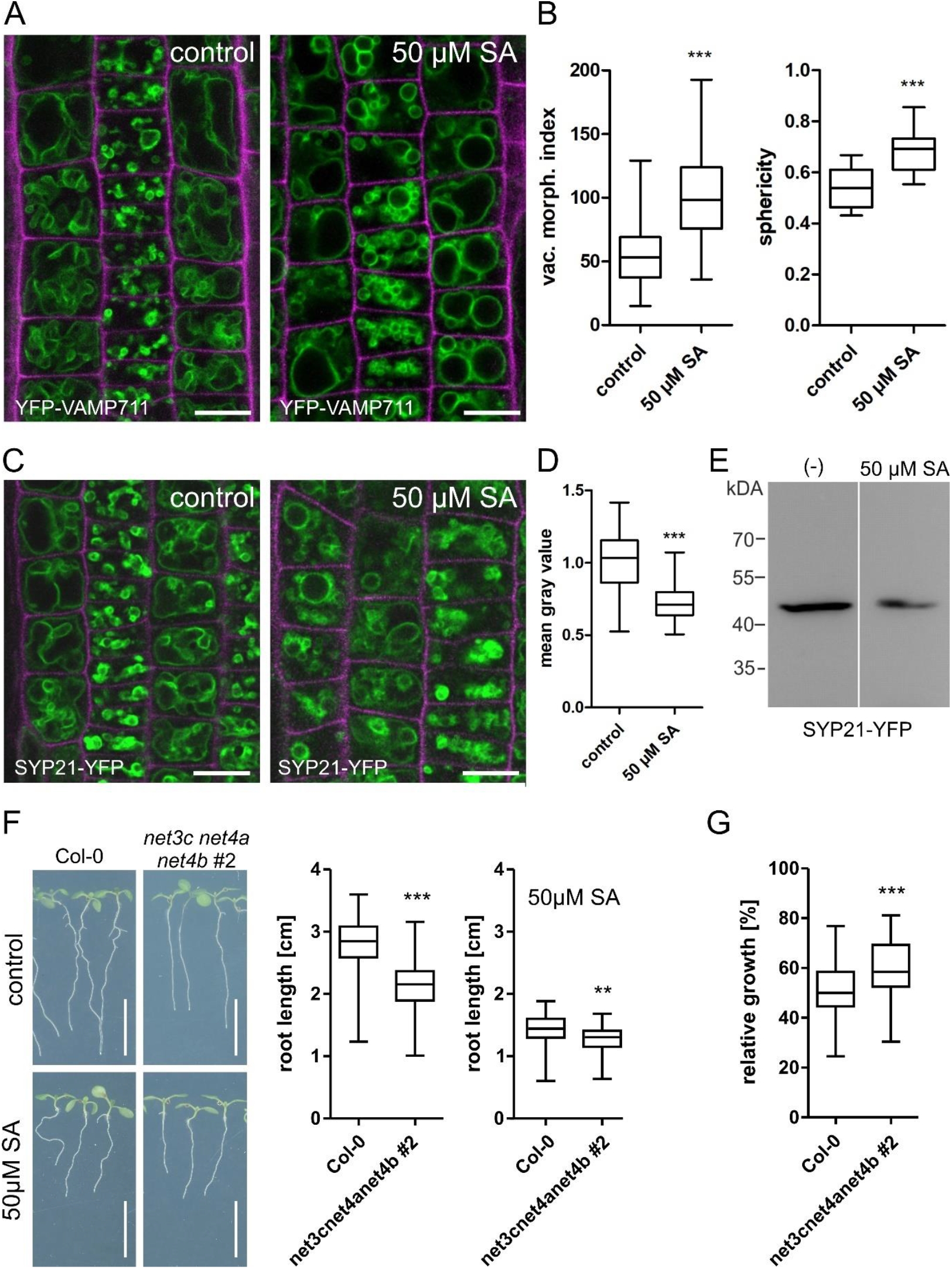
SA rapidly changes vacuolar morphology. A, The tonoplast marker line YFP-VAMP711 was used to visualize vacuole changes after 24h SA treatment (50 µM). Scalebar: 14 µm. B, Vacuolar morphology index (VMI) was calculated as a proxy for vacuole size (n=68). Sphericity was quantified using the Imaris software (Oxford Instruments) on 3D vacuole models (n = 14) shown in Fig. 4. Whisker plot with Student’s t-test. *** p≤0.001. C, Representative images of the SNARE marker line SYP21-YFP before and after 24h SA treatment (50 µM). Scale bar: 15 µm. D, Quantification of SYP21-YFP intensity. SA-treated samples were normalized to the DMSO control (n = 36). Whisker plot with Student’s t-test. *** p≤0.001. E, Western blot detection of SYP21-YFP levels in roots using GFP antibodies. F, Root length of 7-days-old seedlings grown on ½ MS+ agar control plates (n Col-0 = 54, n *net3cnet4anet4b* #2 = 56) and plates supplemented with 50 µM SA (n Col-0 = 56, n net3cnet4anet4b #2 = 51). Whisker plots with Student’s t-test. ** p≤0.01, *** p≤0.001. Scale bar: 1 cm. G, Calculation of relative growth by dividing each seedling’s root length grown on SA supplemented plates with the average root length of the respective seedlings grown on control plates (n Col-0 = 56, n *net3cnet4anet4b* #2 = 51). Values are given as percentages. Whisker plot with Student’s t-test. *** p≤0.001.

### SA application does not Impact Vacuolar Occupancy

Despite the distinct changes of vacuolar morphology after SA treatment, it was still unclear if SA affects vacuolar size. To measure this, we used the luminal vacuole dye BCECF in combination with propidiume iodide, staining the cell wall, for 3D imaging (Scheuring et al., 2015). To this end, we captured z-stacks of root cells from four different positions, including the early -and late meristem, as well as the early and -late elongation zone (Fig. 5A). Relative vacuole size (vacuolar occupancy) was determined by calculating vacuolar volume and corresponding cellular volume based on the respective 3D models. Since it has been shown before that auxin decreased vacuolar occupancy in meristematic cells (Scheuring et al., 2016), it was included as positive control. Remarkably, auxin treatment decreased vacuolar occupancy in cells from all positions, while SA treatment did not change the relative vacuole size at any position (Fig. 5B and 5C). However, SA-inducded changes of vacuolar morphology were still evident in the 3D models (Fig. 5B), indicating that morphology but not the relative size of the vacuole is affected. Notably, when seedlings were grown on SA for 7 days we indeed could observe a reduction of vacuolar occupancy in the early -and late meristem, as well as the early elongation zone (Fig. S4B and C). Decrease of vacuolar occupancy, however, was not as strong as the auxin-induced decrease at all positions.

**Figure 5:**
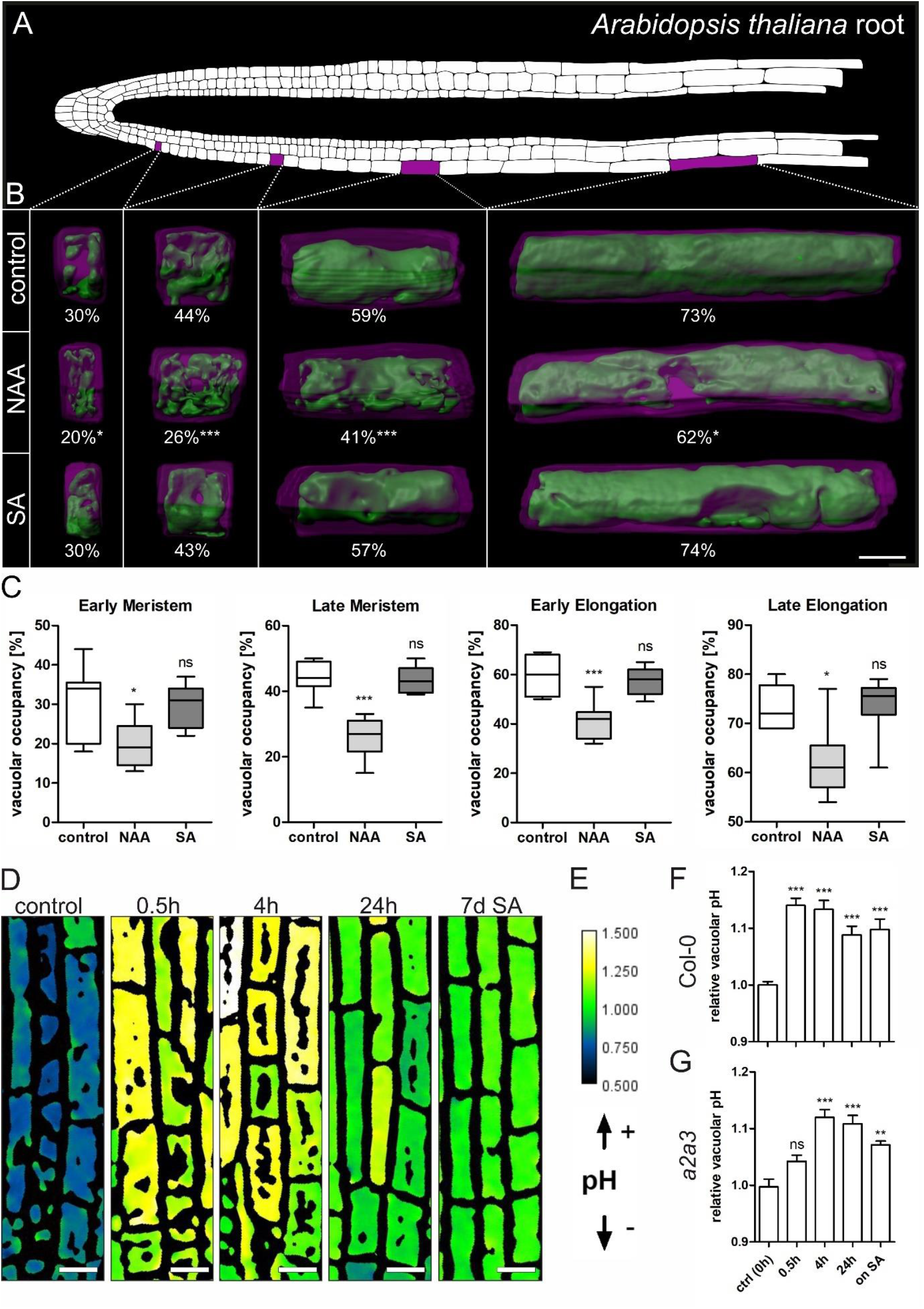
SA induces changes in vacuolar occupancy and increases vacuolar pH. A, Schematic representation of an *Arabidopsis thaliana* root showing epidermis, cortex, and endodermis cell files. The investigated regions are marked in magenta. B, 6-day-old seedlings were grown on ½MS+ medium and transferred for 24h on medium supplemented with 50 µM SA and 250 nM NAA. The seedlings were stained with PI as well as BCECF and z-stacks were acquired. Using these, 3D reconstructions of the vacuole (green) and its corresponding cell (magenta) were generated, and relative vacuole size (vacuolar occupancy) quantified. The different columns are from the left to the right: early meristem, late meristem, early elongation and late elongation zone. The numbers under each picture represent the mean percentage of occupancy of the vacuole. Note: the recess seen in many models indicates the position of the nucleus. Scale bar: 10 µm. C, Quantification of vacuolar occupancy in the four root regions (n = 10). Whisker plot with Student’s t-test. *** p≤0.001, ** p≤0.01, * p≤0.05. D, Col-0 was grown on ½MS+ plates and treated with 50 µM SA for 0.5h, 4h, 24h, and 7 days. Seedlings were stained with BCECF, and ratiometric images were created by dividing the λex 488 intensity by the λex 458 intensity. Scale bar: 15 µm. E, Color code (white to black) reflects the 488/458 intensity (high to low), indicating changes in pH. F, Quantification of vacuolar pH using BCECF, normalized to the DMSO control (n = 44). One-way ANOVA with Tukey post-hoc test. *** p≤0.001.

The general SA-induced changes of the vacuole to a more spherical shape was observed in all experiments and was somewhat reminiscined of the phenotype resulting from the gene knockout of three tonoplast proton pumps: *fugu5-1 vha-a2 vha-a3* (Kriegel et al., 2015). The triple mutant is lacking V-ATPase and V-PPase activity resulting in a significantly higher vacuolar pH. Using the previously employed BCECF in ratiometric measurements for pH determination, we determined pH changes upon exogenous SA application. Surprisingly, already after 30 minutes of SA treatment, a significant increase in vacuolar pH was detected (Fig. 5D-E). This effect lasted for all tested time points, including seedling growth on SA containing media for 7d (Fig. 5F). Since the *fugu5-1 vha-a2 vha-a3* mutant grows very poorly (Kriegel et al., 2015), we used the V-ATPase double mutant *vha-a3-vha-a3* (Krebs et al., 2010) instead to test whether V-ATPase activity might be involved in mediating the SA effect. However, BCECF-based measurements of *vha-a3-vha-a3* showed an intitial delay in pH increase but behaved similarly to the wt at later time points (Fig. 5G).

In tobacco suspension cells it has been observed that glycosylated SA is transported into the vacuole, probably by employing a H^+^-antiport mechanism (Dean et al., 2005). In case such a transport mechanism would exist in Arabidopsis as well, we hypothesized that manipulation of glycosylated SA levels would impact vacuolar pH and eventually growth. To test for this, we used a knockout and an overexpression line of the glucosyltransferases UGT76B1: while there was only little change in vacuolar pH detectable (data not shown), the overexpression line was more resistant than the wt, while *ugt76B1* was significantly more sensitive to SA-induced root growth inhibition (Fig. S5). This could indicate that increased glycosylation modulates the SA response by sequestration of the phytohormone in the vacuole.

## Discussion

Only recently, the function of the phytohormone salicylic acid within plant growth and development came into focus. Previously, many studies described a growth-inhibiting effect by high exogenous or endogenous SA levels (Pokotylo et al., 2022; Rivas-San Vicente and Plasencia, 2011). In some cases, this inhibition was explained by diminished resources due to the classical *growth-defence trade-off*: the plant^’^s energy pool needs to be used for all arising tasks. In extreme situations, e.g. pathogen attack, energy resources must be invested for defence reactions but, ergo, are not sufficient anymore to maintain optimal growth (Huot et al., 2014). More recently, it was proposed that low SA concentrations (<250 µM) in Arabidopsis might have a role in growth and development and only higher concentrations contribute to plant defence (Pasternak et al., 2019). However, we do observe a dosage-dependent inhibition of root growth and already with 50 µM SA a root length reduction of about 50% (Fig. 1). In Arabidopsis and tobacco it has been shown that already relatively low concentrations of SA and its functional analogues INA and BTH induce systemic acquired resistance (SAR) (Ward et al., 1991; Delaney et al., 1994; Lawton et al., 1996) and thus it seems plausible that growth-inhibition and activated plant defence occur simultaneously. For independent regulation, this would imply that sensing and/or signalling of the two processes must be diverging at some point. In line with this, signalling of the *bona fide* SA receptors NPR1, 3 and 4 are not required for the transmission of SA-dependent growth inhibition (Fig. S1) (Tan et al., 2020). This suggests that another signalling pathway, potentially exclusively to regulate growth, might exist.

Another explanation might be the observed auxin-SA crosstalk: lateral root initiation, increased cell division rates and irregular cell file development all correspond to changes of cellular auxin (Pasternak et al., 2019). It has been shown that SA directly binds to a phosphatase (PP2a) which in turn regulates the polarity and thereby activity of the auxin efflux carrier PIN2 (Tan et al., 2020). Remarkably, also other cellular functions, such as endocytosis, are directly affected by SA (Du et al., 2013). This raises the question to which extent SA function requires auxin. Using the R2D2 auxin reporter, we indeed detected strong auxin accumulation in response to SA (24 h) treatment in meristematic root cells (Fig. 2). Additionally, inhibition of cell elongation and loss of cell size differences in tricho-and atrichoblasts upon SA treatment resemble effects induced by exogenous auxin application (Löfke et al., 2013; Löfke et al., 2015a). The findings that the auxin receptor triple mutant *tir1afb2afb3* and the PIN2 mutant *eir1-4* are fully sensitive to SA-induced root inhibition (Fig. 3), however, argue against auxin being solely responsible to conduct SA impact.

The observed SA effect on vacuolar morphology differs massively from the one induced by auxin: while SA induces homotypic vacuole fusion, auxin treatment led to highly constricted vacuoles with decreased relative size (Löfke et al., 2015a; Scheuring et al., 2016). The partial resistance of the *net3cnet4anet4b* triple mutant, which possess fragmented vacuoles, hints towards a correlation between homotypic vacuole fusion and root growth inhibition. On the other hand, the relative size (vacuolar occupancy) of SA treated vacuoles did not decrease questioning the space-filling function of the vacuole to be decisive to regulate growth here (Kaiser et al., 2021; Kaiser and Scheuring, 2020; Krüger and Schumacher, 2018; Scheuring et al., 2016). The observed pH-increase on the other hand, occurred in parallel to the homotypic vacuole fusions within minutes. Interestingly, alkalinization of the V-ATPase double mutant *vha-a2vha-a3* was strongly delayed (Fig. 5), not responding to SA in the first 30 min. When compared to wild type, V-ATPase activity of the *vha-a2 vha-a3* double mutant is severely decreased and vacuolar pH is increased by 0.5 units (Krebs et al., 2010). Thus, one explanation for the observed delay in pH increase might be the higher basic starting pH of the double mutant prior SA treatment. Although not experimentally determined yet, this could indicate that SA affects vacuolar pH by directly inhibiting V-ATPase activity. There is, however, indirect evidence suggesting that SA increases the activity of a plasma membrane H^+^-ATPase in response to salt stress (Jayakannan et al., 2013). Another possible explanation for the SA-induced rapid pH increase might be the need for protons to ensure transport of glycosylated SA into the vacuolar lumen. In tobacco cells, compartmentalization of the glycosylated phytohormone in the vacuole occurs via an H^+^-antiport mechanism which is energized by the proton gradient resulting based on active tonoplast proton pumps (Dean et al., 2005). In Arabidopsis, glycosylated SA has been detected in vacuolar membrane-enriched vesicles and stimulated by MgATP, while vanadate (an ABC transporter inhibitor) and bafilomycin A_1_ (a vacuolar H^+^-ATPase inhibitor) inhibited uptake (Vaca et al., 2017). This suggests that glycosylated SA involves both an ABC transporter and an H^+^-antiporter. Notably, knockout and overexpression of the small-molecule glucosyltransferases UGT76B1 (Saint Paul et al., 2011) resulted in differential sensitivity in respect to SA-induced root growth inhibition (Fig. S5). This could indicate that the measured increase in vacuolar pH is an indirect effect, resulting from the energy-dependent inactivation of SA in the vacuole.

Taken together, SA inhibits growth by limiting cell elongation. These size limitations are accompanied by changes of vacuolar morphology and a rapid pH increase. By interfering with V-ATPase function, SA could directly impacts basic cellular functions such as nutrient storage in the vacuole and general vesicle trafficking (Krebs et al., 2010). The central role for V-ATPases in plant growth and development has been established decades ago (Schumacher et al., 1999). SA-induced growth regulation by targeting the vacuolar proton gradient, however, would represent an entirely novel mechanism.

## Materials & methods

### Plant Material and Growth Conditions

*Arabidopsis thaliana* ecotype Columbia-0 (Col-0) was used as wild type control. The following marker lines were described previously: R2D2 (Liao et al., 2015), PIN2-GFP (Abas et al., 2006) YFP-VAMP711 (Wave 9Y) (Geldner et al., 2009), SYP21-YFP (Robert et al., 2008). The following mutant and transgenic lines were described earlier: *nahG* (Friedrich et al., 1995), *npr1-1 npr3-1 npr4-3* triple mutant (*npr134*) (Zhang et al., 2006), *eir1-4* (Abas et al., 2006), *pp2aa1-6* (*pp2a*) (Blakeslee et al., 2008), *vha-a2 vha-a3* (*a2a3*) (Krebs et al., 2010), UGT76B1^OE^ and *ugt76b1b1* (Saint Paul et al., 2011), *net3cnet4anet4b* (Kaiser et al., 2023). Seeds were surface sterilized with ethanol and plates were stratified at 4 °C for 1–2 days in the dark and grown vertically at 22 °C under a 16 h light/8 h dark-cycle. For seedling growth, half-strength Murashige and Skoog (MS) medium (Duchefa, Netherlands), including 1% (w/v) sucrose (Roth, Germany), 2.5 mM MES (Duchefa) and 1% (w/v) Phytoagar (Duchefa) was used at pH 5.7.

### Chemicals and Treatments

The dyes propidium iodide (PI) and BCECF-AM were acquired from Life Technologies (CA, USA). The synthetic auxin α-Naphthaleneacetic acid (NAA), salicylic acid and its analogues BTH and INA were obtained from Duchefa (Netherlands). Except PI, all chemicals were dissolved in dimethyl sulfoxide (DMSO).

### Confocal Microscopy

Live cell imaging was performed using a Zeiss LSM880, AxioObserver SP7 confocal laser-scanning microscope, equipped with either a Zeiss C-Apochromat 40×/1.2 W AutoCorr M27 water-immersion objective or a Plan-Apochromat 20x/0.8 M27 objective (INST 248/254-1). Vacuole staining was carried out as described previously (Scheuring et al., 2015). Cell walls were stained by mounting roots in 0.01 mg/ml PI solution. Fluorescence signals were acquired for GFP/YFP/BCECF (excitation/emission 488 nm/500–571 nm), PI (excitation/emission 543 nm/580–718 nm), and processed using Zeiss software ZEN 2.3 or Fiji software (https://imagej.net/Fiji). Z-stacks were recorded with a step size of 500 nm (for 3D reconstruction of cells and vacuoles to assess vacuolar occupancy.

### Phenotype Analysis

Estimation of maximal cell length was done as described previously (Löfke et al., 2015a). Root length, calculation of vacuolar morphology index (vac. morph. index, VMI) and vacuolar occupancy of the cell were carried out as described previously (Kaiser et al., 2019).

The meristem length was measured by staining 7-day-old seedlings with propidium iodide (PI) for 20 minutes, followed by image acquisition using a CLSM microscope. The distance between the quiescent center (QC) and the first elongating epidermal cell was quantified and determined as meristem length. For measuring hypocotyl length, seeds were sown on plates and exposed to light conditions for 8 hours to initiate germination. Subsequently, the plates were kept in the dark for an additional 5 days at a constant temperature before documentation. Hypocotyl length was measured using ImageJ by determining the distance from the base of the seedling to the cotyledons. The length of atrichoblasts and trichoblasts was measured by taking CLSM pictures of the epidermal meristem. Here, the length of 3^rd^,4^th^,5^th^ and 6^th^ atrichoblasts, positioned before the first elongated cell (twice as high as wide), were measured and averaged. Subsequently, the number of the corresponding trichoblasts in the adjacent cell file was counted and averaged to the length of the atrichoblasts. The ratio between the averaged lengths of atrichoblasts and trichoblasts was then calculated for comparative analysis. PIN2 polarity was quantified using ImageJ by measuring the intensity of PIN2 fluorescence on the apical and lateral sides of the cells. Measurements were taken from three cells per root. The polarity ratio was calculated by dividing the intensity on the apical side by that on the lateral side. To assess differences between cell types, the polarity ratios of atrichoblasts and trichoblasts were further divided to determine the relative differences in PIN2 distribution between these two cell types.

### GUS staining

GUS staining was performed as described previously (Béziat et al., 2017). In brief: Seven-day-old plate-grown CYCB1;1::GUS seedlings were fixed in 90% acetone for 30 minutes. Following fixation, the seedlings were washed in Na-phosphate buffer for 1 hour and then put into freshly prepared staining buffer. After 4 hours of staining, the seedlings were transferred into an ethanol:acid mixture to remove chlorophyll. The seedlings were then washed for 10 minutes each in 70%, 50%, and 20% ethanol and finally transferred into clearing solution. Images were captured using a Nikon Axiostar Plus microscope and a binocular microscope (Leica M205 FCA).

### Immunolocalization

Whole mount immunolocalization was conducted using six-day-old Arabidopsis seedlings. Fixation was carried out in 4% formaldehyde solved in microtubule stabilization buffer (MTSB; 25 mm PIPES, 2.5 mm EGTA and 2.5 mm MgSO_4_·7H_2_O; pH 6.8) for 40 min. After washing (MTSB) and rehydration, cell wall digestion (2% driselase in MTSB) and membrane permeabilization (3% NP40 in MTSB) were executed. Subsequent washing was followed by blocking (2%BSA in MTSB) for 2h and treatment with a 1:300 diluted primary PIN2 antibody (Abas et al., 2006) over night at 4°C: After extensive washing, a goat anti-rabbit antibody conjugated to PE-CY5.5 (Invitrogen, USA) was applied in 1:400 dilution for minimum 3h. After final washing and mounting, samples were used for microscopy.

### Immunological detection

For immunological detection, samples were separated via sodium dodecylsulfate polyacrylamide gel electrophoresis (SDS–PAGE). Samples were mixed with loading dye, incubated at 95 °C for 5 min and shortly centrifuged prior to loading. Blotting was performed using either a nitrocellulose (GE Healthcare, USA) or a polyvinylidene difluoride (PVDF) membrane (Sigma-Aldrich). Membranes were blocked with 5 % skim milk powder in TBS-T (150 mM NaCl, 10 mM Tris/HCl pH 8.0, 0.1% Tween 20), washed with TBS-T and then incubated with the first antibody solution. After washing the membrane again, it was probed with the second antibody for at least 1 h. Afterwards, the membrane was washed three more times, submerged with ECL solution (ECL Prime Kit; Amersham, UK) and chemiluminescence signals were detected using the ChemiDoc system from BioRad. For detection, a monoclonal anti GFP antibody (1:500; Roche) was used.

### 3D Surface Rendering

3D reconstruction of cells and vacuoles was performed using Imaris 9.1 (Oxford Instruments). For generation of cell models, the manual drawing function (distance) of the surface creation tool was used. Cell borders were marked according to the signals for PI channel on at least every third slice of a z‐stack. Subsequently, a surface representing the whole cell was created based on the markings. This surface was used to mask the BCECF-AM channel by setting the voxels outside the surface to 0. For generation of vacuole models, the masked channel was used to automatically create a surface corresponding to the vacuole (BCECF) signals. Before completion, the displayed model was visually compared to the underlying BCECF signals and potential adjustments carried out using the absolute intensity threshold option. Parameters such as sphericity and volume were extracted for individual 3D models. Vacuolar occupancy was determined by dividing the volume of the created 3D vacuole by the total volume of the corresponding cell. This calculation provided the percentage of vacuolar occupation within the cell.

### pH measurements using BCECF

Vacuolar pH of 7-day-old Col-0 seedlings was determined using the fluorescent cell-permeant dye BCECF. Before loading the dye, seedlings were exposed to 50 µM SA in liquid ½ MS+ medium for varying durations or grown on SA plates for 7 days. For staining, 10 µM BCECF-AM was added to the SA-supplemented liquid medium, and the seedlings were incubated in the dark for 1 hour. Following staining, the seedlings were washed for 10 minutes in fresh liquid ½ MS+ medium. For microscopy, the fluorophore was excited at 488 nm and 458 nm, respectively, and the emission was detected between 530 and 550 nm. The ratio of fluorescence intensity was determined using ImageJ by dividing the values of the 488-nm-excited images by those of the 458-nm-excited images.

### Statistical analysis

All quantitative data was analyzed using the GraphPad Prism 9 software (). The precise statistical method used is given in the respective figure legends.

## Supplemental data

Figure S1: SA reduces root organ growth independently of the NPR-receptors.

Figure S2: SA treatment impacts cell division, leading to altered meristem size.

Figure S3: BTH and INA do not show SA specific phenotypes.

Figure S4: Changes of vacuolar occupancy upon auxin and SA treatment.

Figure S5: Root growth determination of UGT76B1 knockout and overexpression lines.

## Acknowledgements

For providing published material we are grateful to Anton Schäffner, Jiri Friml and Karin Schumacher. We thank Christian Luschnig for providing the PIN2 antibody and the *eir1-4* mutant. We thank Matthias Hahn for critical reading of the manuscript. This work was supported by grants from the *BioComp* research initiative (Rhineland-Palatinate, Germany) and the German research foundation (DFG; SCHE 1836/4-2) to D.S.

## Contributions

J.M. performed most experiments. Y.K., S.K., C.L. and M.K. also carried out experiments. J.M. and D.S. designed the figures and performed statistical analysis. D.S. conceived the study and J.M. and D.S. wrote the manuscript. All authors saw and commented on the manuscript.

## Corresponding author

Correspondence to David Scheuring.

## Competing interests

The authors declare no competing interests.

## Notes

### Competing Interest Statement

The authors have declared no competing interest.

