## Supplemental Data for "Salicylic Acid restricts cell elongation and induces changes of vacuolar morphology and pH"

### Supplemental Information

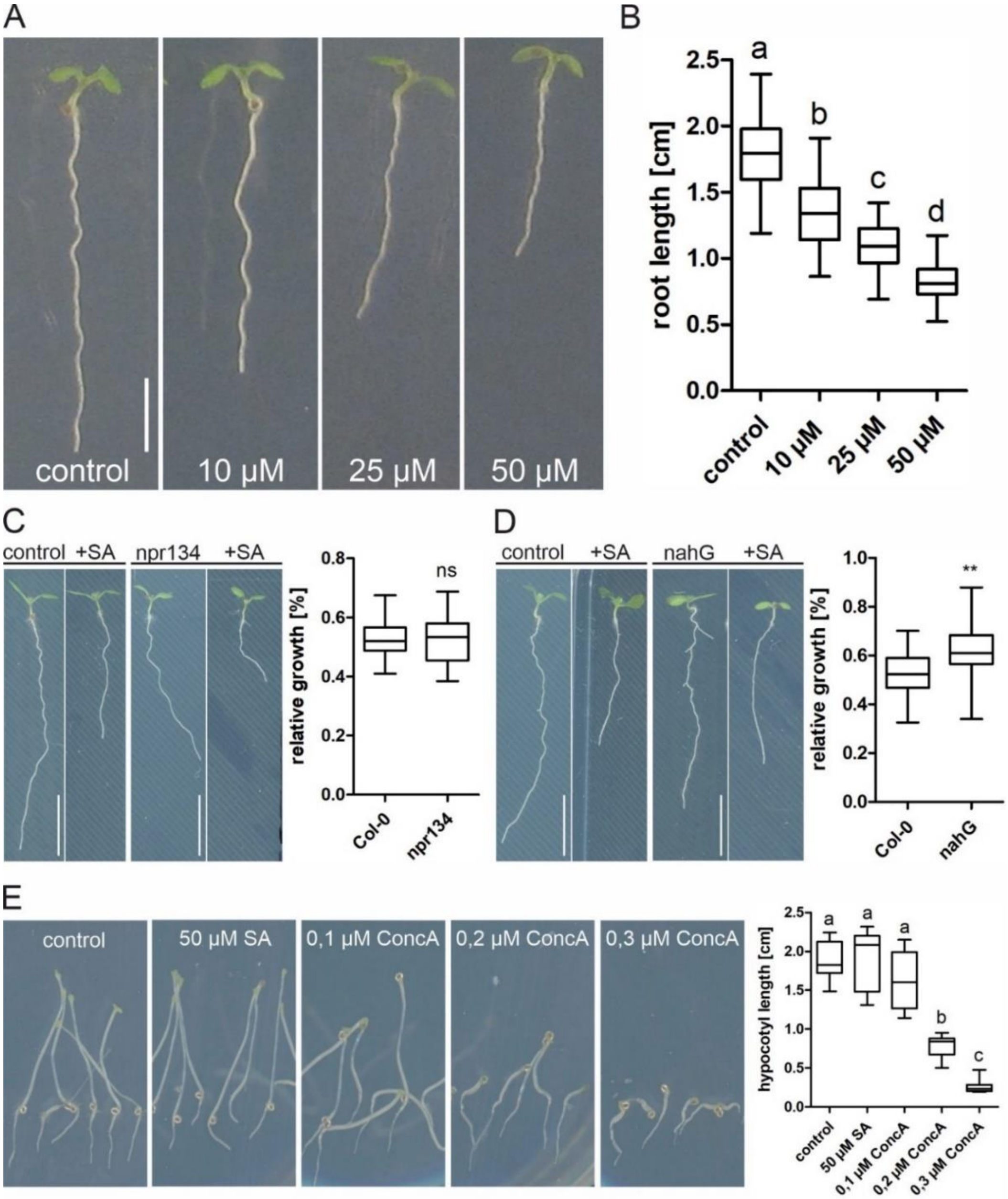


**Figure S1: SA reduces root organ growth independently of the NPR-receptors**. A, Representative images show root length of Col-0 after growing 7 days on different concentrations of SA. Scale bar: 40 mm. Note: values for control and 50 µM SA are included in the graph for Fig. 1D. B, Quantitative analysis of Col-0 root length grown for 7 days on different concentrations of SA. One-way ANOVA test with Tukey post-hoc test. Different letters indicate significant differences with $p \leq 0.001$. C-D, Col-0, *npr134*, and *nahG* seedlings were grown for 7 days on ½MS+ plates with the addition of 50 µM SA. Relative growth was determined by dividing the root length of each seedling grown on SA-supplemented media by the average root length of its corresponding control (n *npr134* = 21, n *nahG* = 20). Student's t-test. ** $p \leq 0.01$. Scale bar: 40 mm. E, Col-0 seeds put on plates with different Concanamycin A concentrations and kept in light for 8 hours after transferring into a dark room for an additional 5 days. Hypocotyl length after 5 days incubation in the dark (n = 10). One-way ANOVA test with Tukey post-hoc test. Different letters indicate significant differences with $p \leq 0.001$.

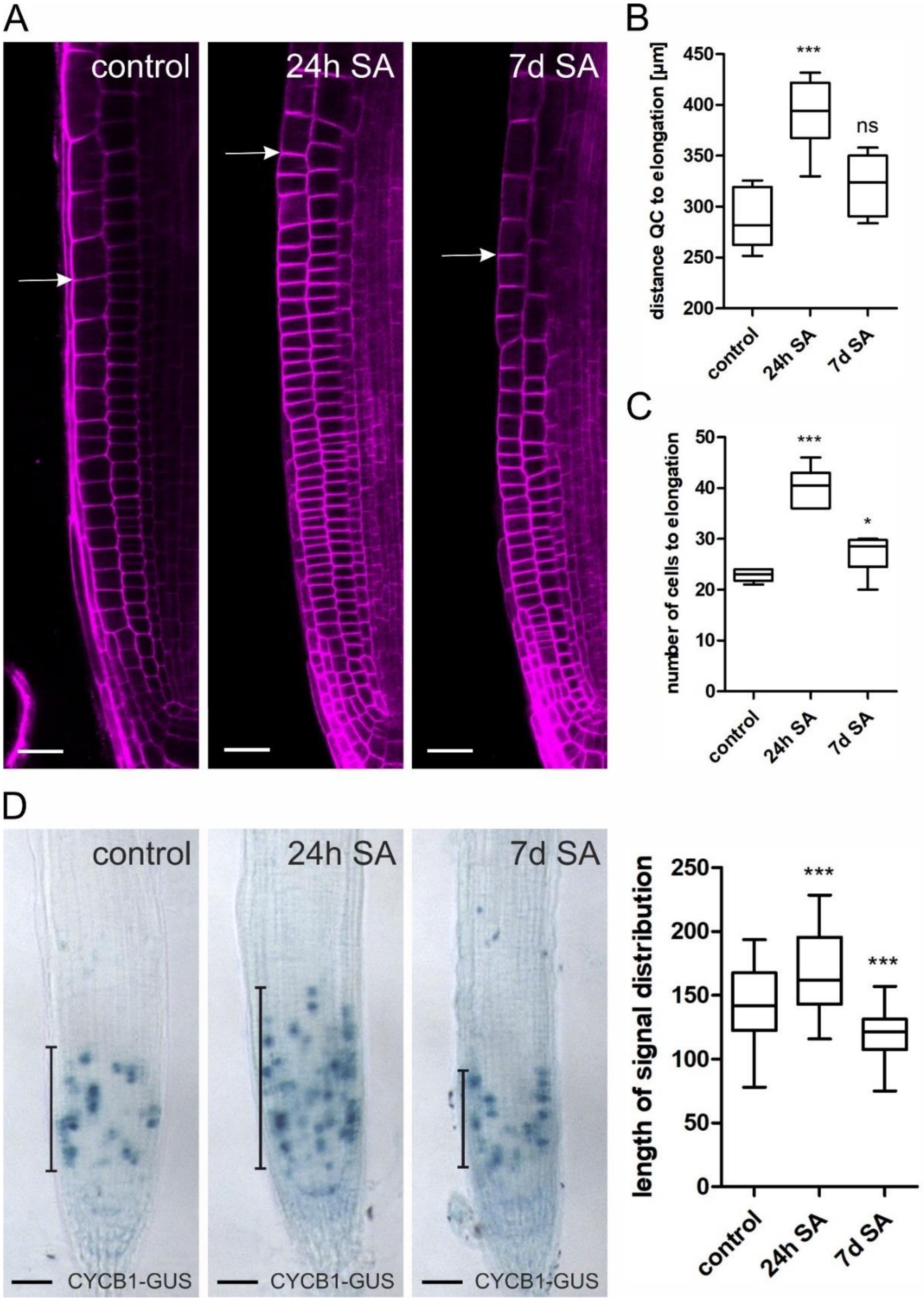


**Figure S2: SA treatment impacts cell division, leading to altered meristem size**. A, Root cross-section stained with PI. The onset of elongation, defined as the first cell being longer than wide, is indicated (white arrow). Scale bar: 30 µm. B, Quantification of the distance of the quiescent centre (QC) to the first elongating cell (n = 8). One-way ANOVA test with Tukey post-hoc test. *** p≤0.001. C, Number of cells from the QC to the first elongating cell (n = 8). One-way ANOVA test with Tukey post-hoc test. * p≤ 0.05, *** p≤0.001. D, The marker for mitotic activity, cyclin B, fused to glucuronidase (*CYCB1::GUS)* was used to follow cell division in the meristem after 24h and 7d SA (50 µM) treatment. The length of signal distribution, indicated by the black bars, was quantified (n = 38). Whisker plot with Student's t-test. *** p≤0.001.

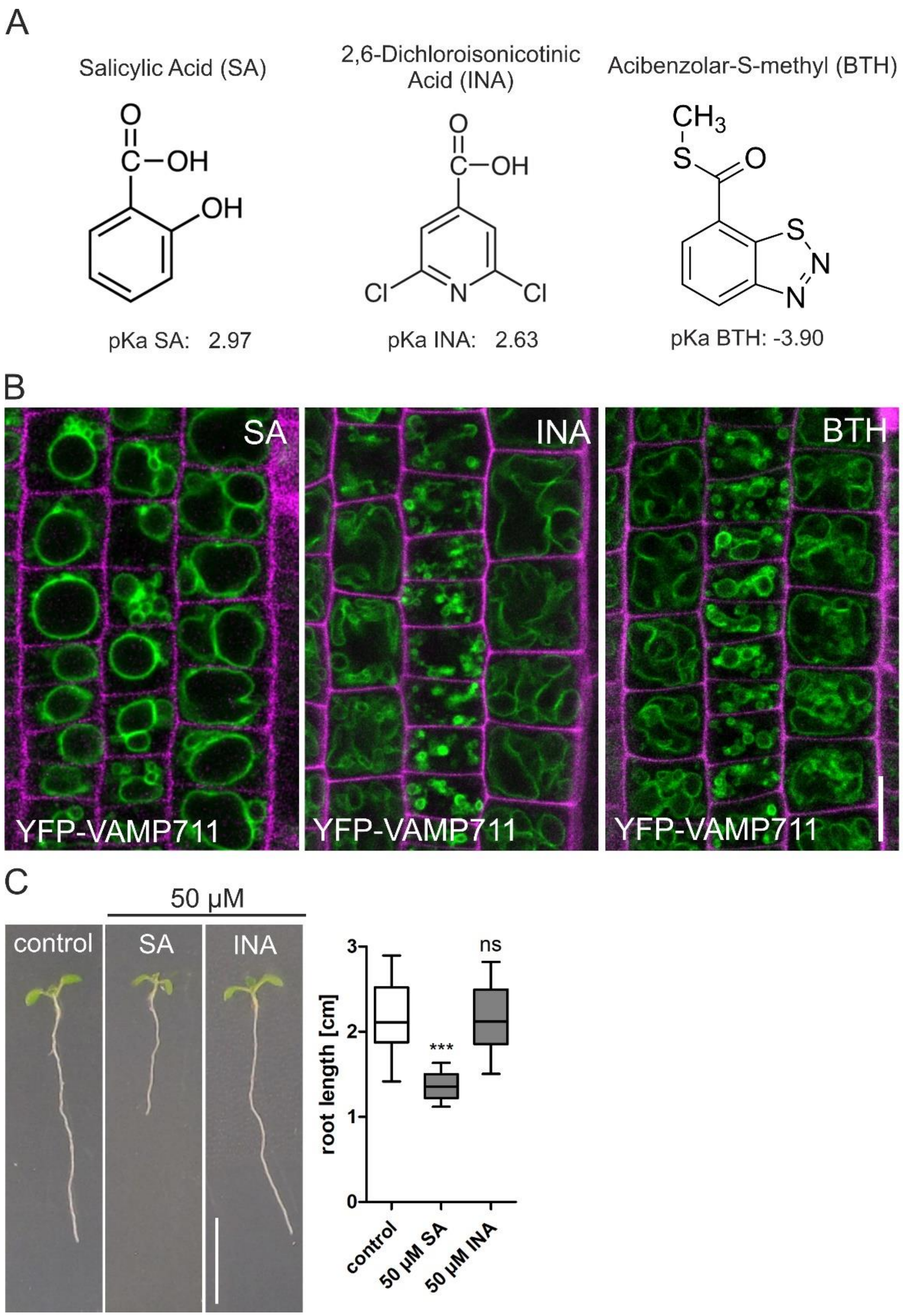


**Figure S3: The functional SA analogs INA and BTH do not show SA-related phenotypes.** A, Structural formula and pKa value of SA, 2,6-dichloronicotinic acid (INA) and acibenzolar-S-methyl (BTH). The tonoplast marker line pUBQ10::YFP-VAMP711 was used to visualize vacuole changes after 24h SA (50 µM), INA (50 µM) or BTH (50 µM) treatment. Scale bar: 15 µm. C, Root growth determination: Col-0 seedlings were grown for 7 days on ½MS+ plates and supplemented with 50 µM SA and INA respectively. The relative growth was calculated by dividing each root length of seedlings grown on SA supplemented media with the average root length of its respective control (n = x). Student's t-test. *** p≤0.001. Scale bar: 40 mm.

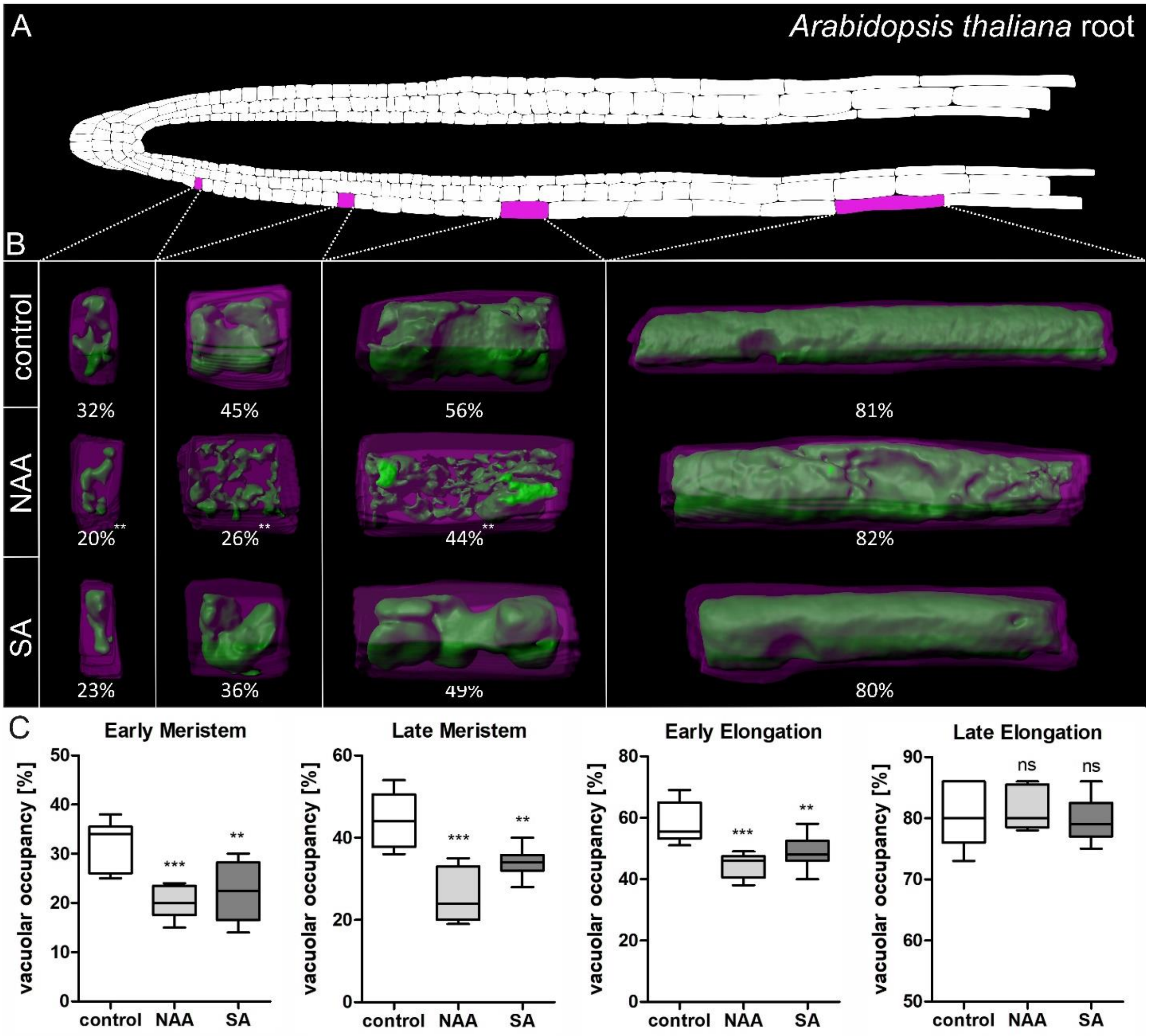


**Figure S4: Changes of vacuolar occupancy upon auxin and SA treatment.** A, Schematic representation of an *Arabidopsis thaliana* root showing epidermis, cortex and endodermis cell files. To visualise the investigated region, the corresponding cells were marked in magenta. B, Col-0 seedlings were grown for 7 days on ½MS+ medium supplemented with 50 µM SA and 250 nM NAA. The seedlings were stained with PI as well as BCECF and Z-stacks with a CLSM were acquired. With these, 3D reconstructions of the vacuole (green) and its corresponding cell (magenta) were generated, and the vacuolar occupancy quantified. The different columns are from the left to the right: early meristem, late meristem, early elongation and late elongation zone. The numbers under each picture represent the mean percentage of occupancy of the vacuole. C, Quantification of the vacuolar occupancy for the four different root regions (n = 10). Whisker plot with Student's t-test. *** $p \leq 0.001$ and ** $p \leq 0.01$.

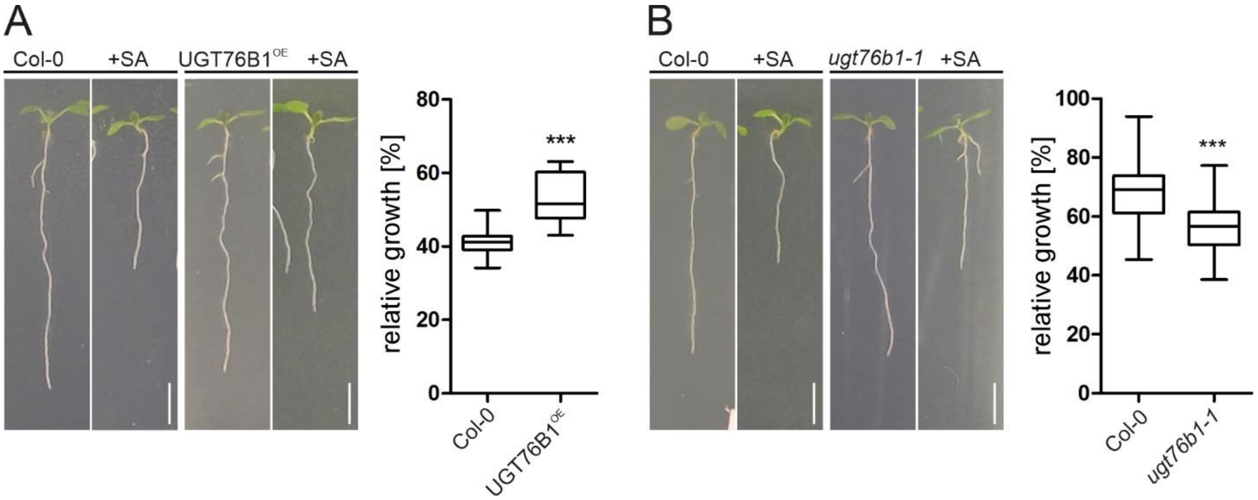


**Figure S5: Root growth determination of UGT76B1 overexpression and knockout lines upon SA treatment.** A-B, Col-0, UGT76B1$^{OE}$, and *ugt76b1-1* seedlings were grown for 7 days on ½ MS+ plates supplemented with 50 µM SA. The relative growth was calculated by dividing the root length of seedlings grown on SA supplemented media with the average root length of its respective control (n Col-0 = 17, n UGT76B1$^{OE}$ = 19 for A and n Col-0 = 43, n *ugt76b1-1* = 42). Whisker plot with Student's t-test. *** $p \leq 0.001$. Scale bar: 30 mm.
